# Dissect mitochondrial activity by transcriptome data with mitology R package

**DOI:** 10.1101/2024.11.27.625451

**Authors:** Stefania Pirrotta, Massimo Bonora, Enrica Calura

## Abstract

Mitochondria are dynamic organelles that play crucial roles in energy transformation, biosynthesis, and cellular signaling. They actively process biological information, detecting and reacting to both internal and external stimuli. Through intricate physical interactions and diffusion mechanisms within cellular networks, mitochondria integrate diverse inputs and generate signals that finely adjust cellular functions and overall physiology. As a result, the phenotypic expressions of impaired mitochondrial function can exhibit high variability. High-throughput transcriptomic data can capture these changes, but traditional pathway analyses performed on common databases often struggle to specifically detect mitochondrial alterations. This is primarily due to the size of pathways, where the mitochondrial component is typically a small portion of the examined signaling network. To allow a specific exploration of mitochondrial activity through transcriptomic profiles, we developed the mitology R package. We started with a collection of genes whose proteins localize in to the mitochondria, derived from specialized databases like MitoCarta, IMPI, MSeqDR, and from the Gene Ontology database (genes annotated with terms related to ‘mitochondri-’ in their description). Then, exploiting the mitochondrial gene list, mitology provides ready-to-use implementations of MitoCarta pathways and also a reorganization of general pathway databases, like Reactome and Gene Ontology, with a specific focus on the pathways related to mitochondria activity. Finally, mitology utilizes these mitochondria-focused pathways to conduct mitochondrial pathway analyses, enabling single-sample assessments. Furthermore, we extended the functionality of our package to accommodate classical gene expression data, as well as the newest techniques of single-cell and spatial transcriptomics. Mitology emerges as a novel R package adept at dissecting and unraveling mitochondrial activity, serving as an instrument for conducting targeted mitochondrial studies.

## Introduction

Mitochondria are cellular organelles whose main role is related to the aerobic respiration to supply energy to the cell. In addition to this, they are also involved in other tasks, such as signaling, cellular differentiation, and cell death, as well as maintaining control of the cell cycle and cell growth [1]. For this reason, the dysfunction of mitochondria activity has been implicated in several human disorders and conditions, such as mitochondrial diseases, cardiac dysfunction, heart failure [2] and autism [3]. Moreover, mitochondria are involved in cancer diseases, as they have a role in initiation, progression and cancer metastasis formation. Studying mitochondrial behavior in health and disease conditions is opening new possibilities of understanding disease mechanisms and providing new treatments. Indeed, the possibility to target mitochondria for cancer therapies is increasingly becoming a reality in biomedical research [4].

The analysis of high resolution transcriptomic data can help in better understanding mitochondrial activity in relation to the gene expression dynamics. Even if a curated mitochondrial specific pathway resource exists (MitoCarta3.0, which provides a list of mitochondrial genes organized into mitochondrial pathways [5]) computational tools for mitochondrial-focused pathway analysis are still lacking.

In the field of gene set analysis, general purpose resources are pathway databases, such as Reactome [6] and the Kyoto Encyclopedia of Genes and Genomes (KEGG) [7], and the Gene Ontology (GO) project [8], which aim at providing gene set categories for high-throughput data analysis for all the cellular contexts. In the vast majority of pathways contained in these public databases, the mitochondrial-specific component of cellular signaling constitutes only a small portion, which is often overshadowed by the non-mitochondrial signaling. And, even if the information on protein localization is present, pathway analysis typically does not account for whether the activated signal involves the mitochondrion or not.

Thus, we decided that a tool able to focus specifically on the mitochondrial part of pathways could allow tailored and more specific analysis on this interesting organelle and we started to develop mitology to provide actionable and fast mitochondrial phenodata interpretation of high-throughput data. In its present form, mitology provides mitochondrial pathways and ways to perform and visualize mitochondrial pathway analyses.

The development of mitology can be described as multiple phases: (i) collection of mitochondrial genes; (ii) organization of mitochondrial genes in gene sets; (iii) definition of utilities for pathway analyses in multiple gene expression omics; (iv) development of graphical tools for results visualization.

## Materials and Methods

### The collection of mitochondrial genes

We collected the mitochondrial genes from multiple resources: MitoCarta 3.0 [5], IMPI [9], MSeqDR [10] and GO [8]. We also considered the use of pathway databases, such as Reactome or KEGG, but no mitochondrial-specific pathways are described - pathways with “mitochondri-” substring in the title - even if multiple mitochondrial genes are annotated. Genes collected in the MitoCarta3.0 database were downloaded from the site (https://www.broadinstitute.org/mitocarta/mitocarta30-inventory-mammalian-mitochondrial-proteins-and-pathways) for the human species (“Human.MitoCarta3.0.xls” file) curated by the Broad Institute. Genes collected in the IMPI database were downloaded from the site (https://www.mrc-mbu.cam.ac.uk/research-resources-and-facilities/impi) curated by the University of Cambridge. The fourth and last version of the database was selected (“IMPI-2021-Q4pre”). From this collection, we kept the genes encoding verified mitochondrial proteins with “gold standard” evidence of mitochondrial localisation; the ones encoding associated mitochondrial proteins with evidence of mitochondrial localisation, but lacking visual confirmation; and the genes encoding ancillary mitochondrial proteins with no evidence of mitochondrial localisation, but reported to affect mitochondrial function or morphology. Genes collected in the MSeqDR database were downloaded from the MSeqDR site (https://mseqdr.org/). To retrieve the genes from GO, we selected the terms from the Cellular Component category with “mitochondri-” substring in the GO term description. Terms were accessed with the *GO*.*db* R package (v3.20.0). The four lists of genes were unified in a final list of mitochondrial genes.

### The collection of mitochondrial pathways

The whole and original structure from the MitoCarta3.0 database was kept and included in mitology organized in a three-tier hierarchy (Figure 2B).

**Figure 1.**
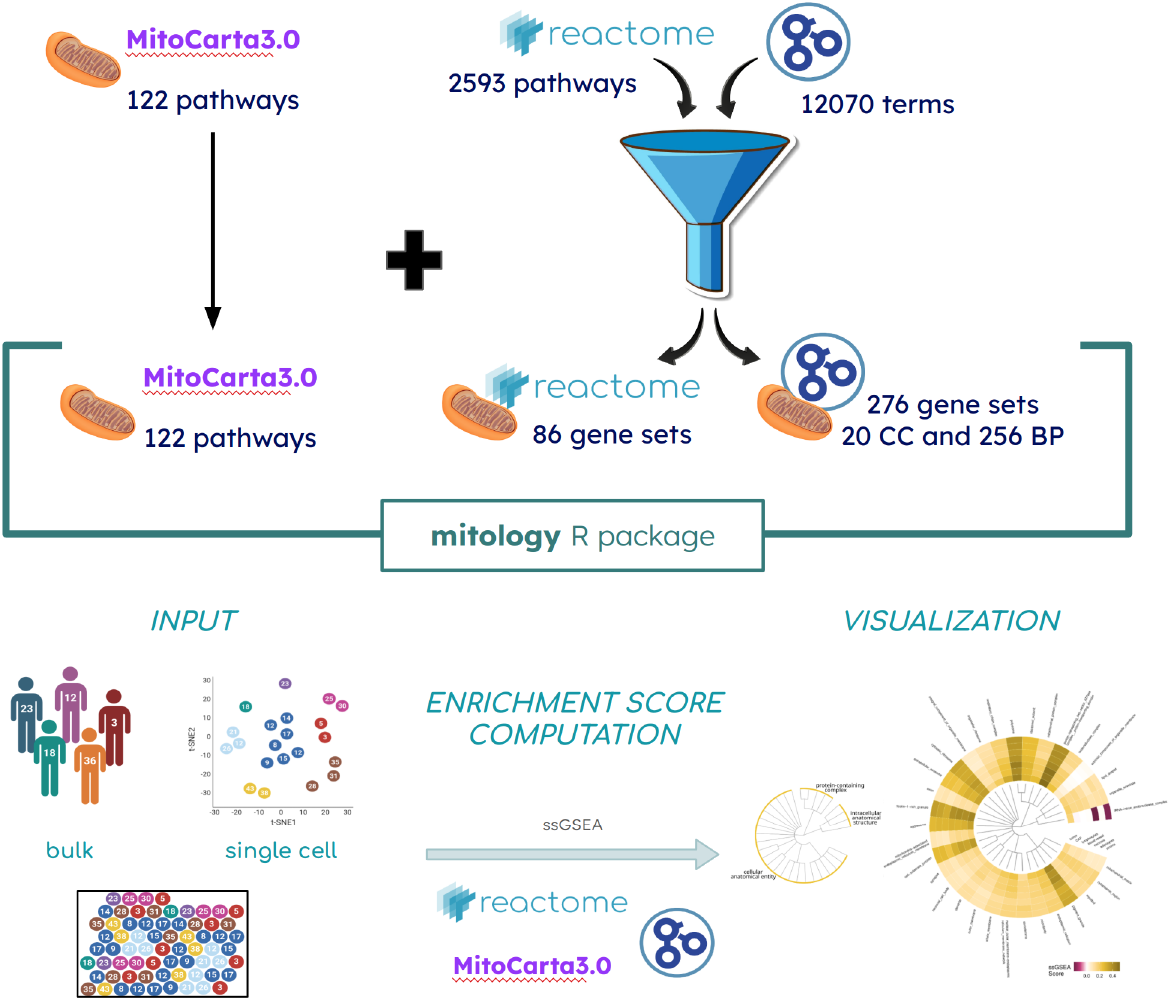
The workflow of the mitology implementation and analysis.

**Figure 2.**
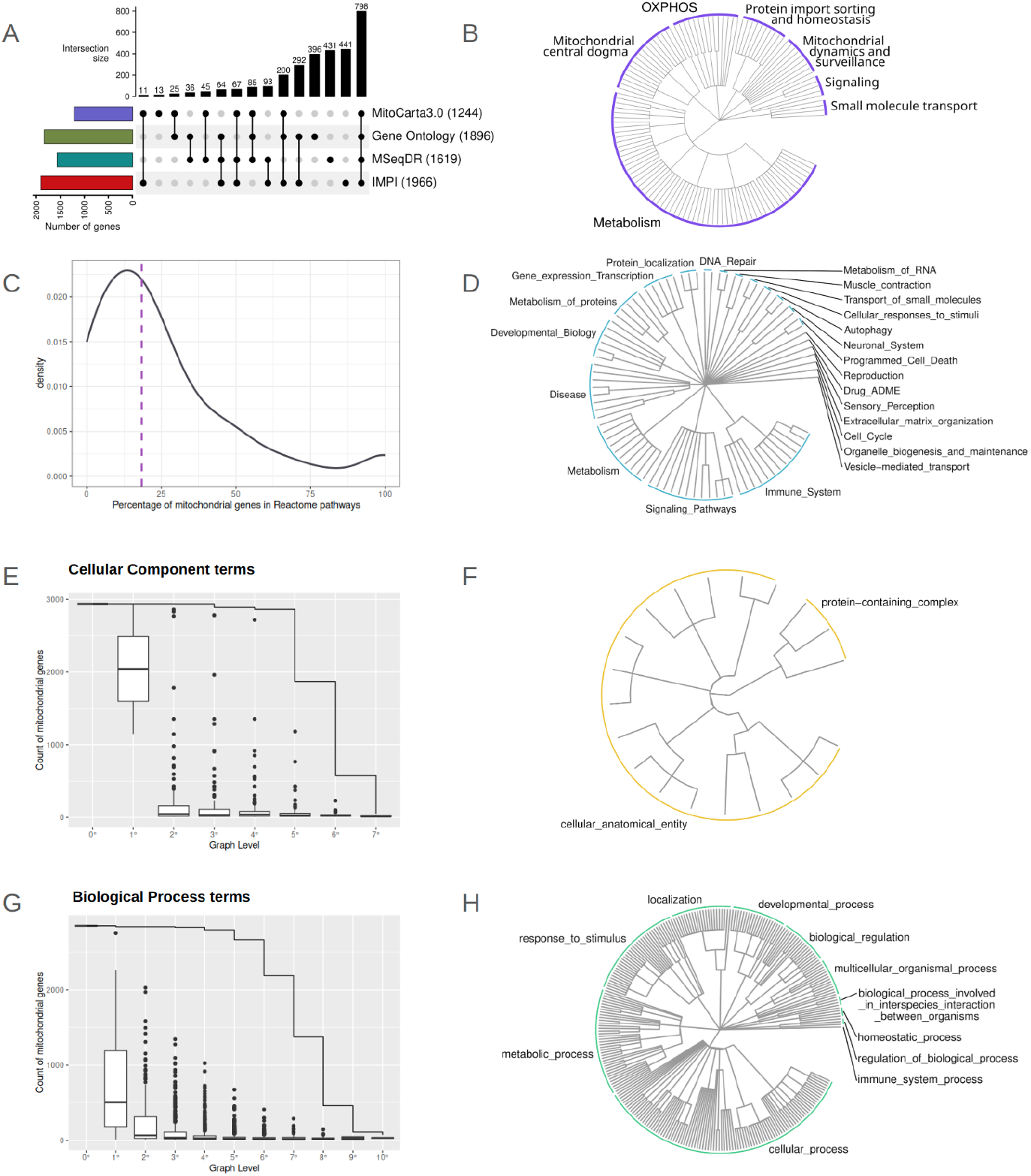
**A** - Upset plot of the intersection between the different four lists of mitochondrial genes. **B** - Circular tree of the MitoCarta3.0 pathways structure. **C** - Density distribution of the percentage of mitochondrial genes inside the Reactome pathways. Median value is indicated by the purple dashed line and is at 18.4%. **E, G** - Boxplots of the number of mitochondrial genes present in each term by term level for GO-CC and GO-BP. **D, F, H** - Circular trees of the Reactome, GO-CC and GO-BP gene sets structures.

In addition to MitoCarta3.0, we decided to propose three more comprehensive mitochondrial-oriented gene set resources exploiting Reactome, GO Biological Processes (GO-BP), and GO Cellular Components (GO-CC).

Starting from the hierarchical organization of GO and Reactome, we applied a gene set step selection to optimize the choice of the most representative mitochondrial gene sets.

We explored the GO-CC and GO-BP terms from the *GO*.*db* R package (v3.20.0). To filter the GO terms we performed a four-steps selection. Firstly, terms were filtered by size, keeping all the ones with at least 10 mitochondrial genes. Then, we pruned the GO trees of CC and BP by the graph levels, filtering out the 0°, 1°, 2° and 3° levels and keeping the terms from the 4° level. Then, we tested the enrichment of the remaining sets for our mitochondrial gene list with the *enrichGO* function from the *clusterProfiler* R package (v4.14.3). Only sets with FDR lower than 0.05 have been passed to the next step. Finally, the last selection was topological, we exploited the GO hierarchical organization selecting the more general enriched gene sets filtering out their offspring terms.

The Reactome pathways were retrieved with the *graphite* R package (v1.52.0). Reactome has a simpler structure with far less level of nested pathways, thus we applied only three filtering steps: number of mitochondrial genes over 10; FDR under 0.05 (enrichment computed with the *phyper* function) and filtering of the offspring pathways.

### Mitology package implementation details

Once we obtained the final terms and pathways respectively from GO and Reactome, we extracted the mitochondrial gene sets by keeping only the genes included in the mitochondrial list. The same names of the original terms/pathways were kept for the gene sets included in mitology. The final gene sets and the corresponding tree-structures of the four databases were kept and included in the mitology package.

Mitology is an open-source R package, available from the GitHub repository (https://github.com/CaluraLab/mitology). It is released under the AGPL-3 license and requires R version 4.2.0 or higher.

## Results And Discussion

Figure 1 graphically outlines mitology development and the workflow of analyses.

The proposed package allows users to compute pathway analysis specifically on mitochondrial gene sets derived from MitoCarta3.0, and on a selection of the Reactome, GO-CC and GO-BP gene sets containing only the mitochondrial genes. Mitochondrial genes reported in mitology are defined as genes that are either part of the mitochondrial genome, part of the mitochondrial proteome, or mitochondrial ancillary proteins. These ancillary proteins may not be located in the mitochondria but are reported to influence mitochondrial function or morphology.

At the time of writing, mitology provides pathway analysis by using the ssGSEA method [11], of all the three types of expression datasets – bulk, single-cell, and spatial transcriptomics data. Additionally it provides utilities for the graphical representation of the enrichment scores: circular heatmaps over the dendrogram of mitochondrial gene set hierarchical structure. Mitology is an open-source R package, available on GitHub (https://github.com/CaluraLab/mitology).

### Comprehensive screening of mitochondrial genes

The initial task was collecting all the mitochondrial genes. Mitochondrial DNA encodes only 13 structural genes, which contribute to the respiratory chain. These include seven subunits of NADH-coenzyme Q reductase (Complex I), three subunits of cytochrome c oxidase (Complex IV), mitochondrial cytochrome b (a component of Complex III), and two subunits of mitochondrial ATP synthase[12]. The vast majority of mitochondrial proteins, however, are encoded by nuclear DNA[13].

Collections of mitochondrial proteins, i.e. all those proteins that localize at mitochondria and/or that directly affect mitochondrial activities, exist. To retrieve these types of genes, we took advantage of public resources that were developed in the last years to define the mitochondrial proteome with the aim to study mitochondrial functions, dysfunctions and diseases. Specifically, we collected mitochondrial genes from the following sources: MitoCarta3.0 [5], the Integrated Mitochondrial Protein Index (IMPI) [9], the Mitochondrial Disease Sequence Data Resource (MSeqDR) [10] and The Gene Ontology (GO) [8].

MitoCarta is a comprehensive catalog of genes encoding the mitochondrial proteome. Its latest version, MitoCarta3.0 [5], includes 1244 manually curated human genes whose proteins are strongly supported to localize to mitochondria. Additionally, it provides annotations linked to a custom ontology comprising 149 mitochondrial pathways.

IMPI is the second curated collection of human genes encoding mitochondrial proteins [9]. The latest version of IMPI includes 1966 genes divided in three groups: (i) 1357 genes encoding verified mitochondrial proteins with “gold standard” evidence of mitochondrial localisation; (ii) 127 encoding associated mitochondrial proteins with evidence of mitochondrial localisation, but lacking visual confirmation; (iii) 482 encoding ancillary mitochondrial proteins with no evidence of mitochondrial localisation, but reported to affect mitochondrial function or morphology.

MSeqDR - a global grass-roots consortium facilitated by the United Mitochondrial Disease Foundation [10] - is the third resource considered. The latest version includes 1619 nuclear and mitochondrial genes linked to mitochondrial biology and disease. Specifically, these genes include all genes from mitochondrial DNA, nuclear genes known to cause mitochondrial disease, nuclear genes known to encode mitochondrial proteins, and genes included on clinical panels or research investigations of mitochondrial biology and disease.

Finally, we took into consideration the GO–CC domain and we extracted all terms containing the substring “mitochondri-” in their descriptions. This approach identified a total of 1896 genes associated with these terms.

At the end, 2996 mitochondrial genes were obtained by joining the four lists. We also considered the use of pathway databases, such as Reactome and KEGG, to collect mitochondrial genes, but very few mitochondrial-specific pathways are described in them - pathways with “mitochondri-” substring in the title - even though multiple mitochondrial genes are annotated in these databases. The numbers and overlaps among these gene lists are visually represented in Figure 2A. Of the 2996 total mitochondrial genes, 57.2% appear in at least two of the four sources. Notably, 798 genes (26.6% of all mitochondrial genes) are shared across all four resources. In contrast, 42.8% of the genes are specific to a single database: 13.2% are unique to GO-CC, 14.4% to MSeqDR, and 14.7% to IMPI. Genes from the MitoCarta3.0 collection, however, show the highest overlap with the other lists and just 13 specific genes.

### Organization of mitochondrial genes in gene sets

Once obtained the mitochondrial gene list, the following task was the selection of the mitochondrial gene sets.

#### MitoCarta3.0

MitoCarta3.0 already provides a comprehensive list of mitochondrial biological processes, which includes 1126 genes manually annotated into seven macro-groups of mitochondrial processes (Metabolism, Mitochondrial central dogma, Mitochondrial dynamics and surveillance, OXPHOS, Protein import, sorting and homeostasis, Signaling, Small molecule transport) further subdivided in 122 smaller subpathways [5].

The whole and original structure from the MitoCarta3.0 database was kept and included in mitology organized in a three-tier hierarchy (Figure 2B). As far as we know, this is the first time that the database is computationally implemented for its use with the R programming language. MitoCarta3.0 is a useful reference for researchers working on mitochondria, but only the 35% of our mitochondrial gene list and the 85% of the MitoCarta3.0 gene list is annotated in its pathways. We believed that the information from the MitoCarta3.0 pathways could be effectively integrated with other gene set resources, such as Reactome and GO, to enhance mitology utilities.

#### Reactome and Gene Ontology

In addition to MitoCarta3.0, we developed three additional comprehensive mitochondrial-focused gene set resources, leveraging Reactome, GO-BP, and GO-CC. In their original annotations, these resources include mitochondrial genes across many pathways and GO terms. Specifically, 1955 out of 2593 Reactome pathways, and 10799 out of 26467 GO-BP terms, as well as 1271 out of 4022 GO-CC terms, contain at least one mitochondrial gene. However, as shown in Figure 2C, the mitochondrial component is minimally represented in the vast majority of these cases.

Starting from the hierarchical organization of GO and Reactome, we applied a gene set step selection to optimize the choice of the most representative mitochondrial gene sets. The scope of the procedure was having the minimum number of gene sets able to classify the maximum number of mitochondrial genes, maintaining biological information as specific as possible, and the gene set size compatible with the analysis, avoiding as much as possible the gene redundancy across sets. Ideally, the gene set size should be sufficiently large to allow robust statistical analyses but sufficiently small to guarantee biological specificity of the term.

GO has a hierarchical structure with multiple levels and highly interconnected terms. To filter the mitochondrial GO terms, we followed a four-step selection process, outlined below. First, we filtered the GO gene set by size, retaining only those terms with at least 10 mitochondrial genes. This step significantly reduced the number of terms studied, while having minimal impact on the number of mitochondrial genes annotated within those terms. Next, we focused on pruning the GO hierarchy to avoid overly broad terms. Figures 2E and 2G display the number of mitochondrial genes in each term by level for CC and BP. The top line shows the cumulative number of genes. In both CC and BP, we observed that up to the fourth level, the total number of genes accounted for more than 95% of the maximum possible number for that category (level 0°). However, from the fifth level onwards, the number of genes began to decrease. Based on this, we pruned the 0°, 1°, 2°, and 3° levels and retained only the terms from the 4° level onward. Then, we tested the enrichment of the remaining sets for our mitochondrial gene list. Only sets with FDR lower than 0.05 have been passed to the next step. Finally, the last selection was topological based on the hierarchical relation across terms: in case of closely related terms we selected the more general enriched gene sets filtering out all the offspring enriched terms.

The filtering procedure for Reactome, which has a simpler pathway hierarchy with fewer levels and pathways, consisted of three steps: i) retaining only pathways with more than 10 mitochondrial genes, ii) applying a false discovery rate (FDR) threshold of 0.05, and iii) filtering out offspring pathways (Figure 2 C, D and Table 1).

**Table 1.** number of terms/pathways after each filtering step.

| | All | number of mito<br>genes $\geq 10$ | graph level $\geq 4^\circ$ | enrichment p<br>value $\leq 0.05$ | remove<br>offspring |
| --- | --- | --- | --- | --- | --- |
| **GO-CC** | 1271 | 418 | 198 | 82 | 20 |
| **GO-BP** | 10799 | 3180 | 2309 | 1292 | 256 |
| **Reactome** | 1955 | 642 | - | 245 | 86 |

After the filtering procedure, the remaining pathways and terms are organized into three trees, shown in Figures 2D, 2F, and 2H. These circular trees represent the selected mitochondrial gene sets from Reactome pathways, GO-CC terms, and GO-BP terms. The resulting graphs preserve the original hierarchical structure, which is also taken into account in the proposed pathway analyses.

Finally, the selected gene sets from Reactome and GO were refined by retaining only the mitochondrial genes.

#### Comparison across mitology gene sets

Once we obtained the final gene sets from the Reactome and GO databases we were interested in seeing how much the MitoCarta3.0 genes overlap with the gene sets from Reactome and GO.

Figure 3A shows the intersection of mitochondrial genes mapped in the gene sets (2950) of each of the considered resources. All but 4 genes from the MitoCarta3.0 pathways are also mapped in at least another gene set from a different resource, while 1898 (64%) of the mitochondrial genes are found only in the gene sets derived from Reactome pathways and GO terms demonstrating the potential complementarity of all the four resources.

**Figure 3.**
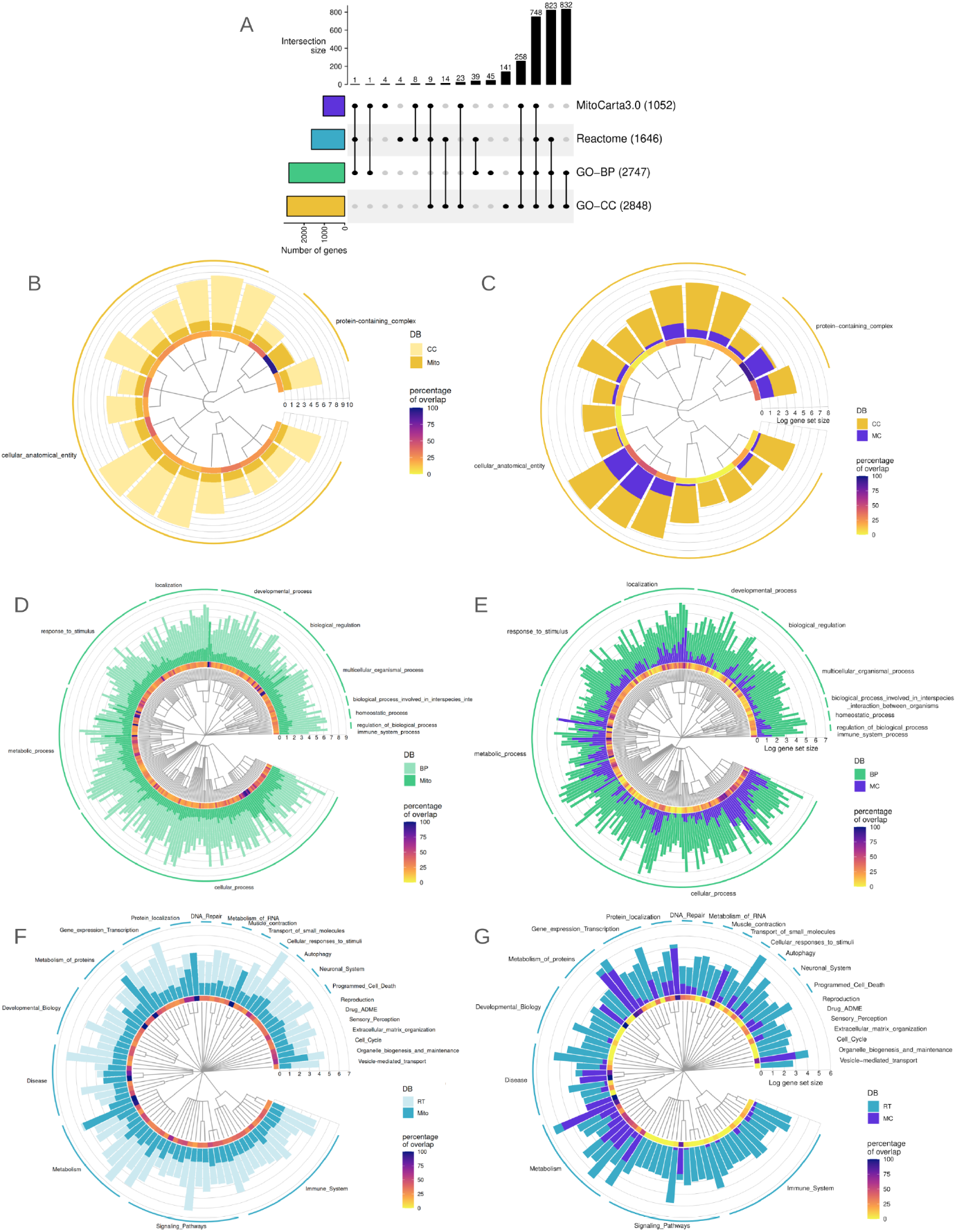
**A** - Upset plot of the intersection of genes included in the four databases of gene sets. The trees of the mitochondrial GO-CC (**B**), GO-BP (**D**) and Reactome (**F**) gene sets, along with the histogram depicting the original pathway size and its respective size after the mitology filtering. The trees of the mitochondrial GO-CC (**C**), GO-BP (**E**) and Reactome (**G**) gene sets, along with the histogram depicting the mitology gene set size and the portion of MitoCarta3.0 genes (in violet).

Figures 3B, 3D, and 3F show, for the selected gene sets of each resource, the size of the original pathway and the size of the pathway in mitology containing only mitochondrial genes. Figures 3C, 3E, and 3G display, for the same selected gene sets, the number of MitoCarta3.0 genes included in each mitology gene set. From the overview provided by these plots, it is clear that, although MitoCarta3.0 pathways are a well-known, manually curated resource dedicated to mitochondria, the other resources, if appropriately filtered, can also be valuable in expanding the range of pathways and gene sets containing mitochondrial genes.

#### Mitochondrial Genes of unknown function

Even when using GO and Reactome, there is a portion of our mitochondrial gene list that remains unannotated in these sets. Specifically, 1350 genes (45% of the total mitochondrial list) are not annotated in Reactome, 266 genes (9% of the total mitochondrial list) are not annotated in GO-BP, and 168 genes (6% of the total mitochondrial list) are not annotated in GO-CC. In total, 46 genes (1.5% of the mitochondrial list) are not annotated in any of the four resources considered. We refer to these as orphan genes, i.e., genes with unknown function.

## Conclusions

In this study, we introduced mitology, a comprehensive resource that provides mitochondrial genes and gene sets, facilitating mitochondria-oriented pathway analysis. The included gene sets are derived from MitoCarta, as well as those filtered from Reactome GO. Our findings highlight that, while MitoCarta3.0 remains a well-established resource for mitochondrial genes, other resources like Reactome and GO - when appropriately adapted - can significantly enhance the annotation of mitochondrial pathways and functions.

Mitology preserves the hierarchical organization of pathways, allowing for more interpretable results. It offers two levels of specificity in gene set analysis: one focused on smaller, specific pathways, and another on broader, parent macro-categories. This structure enables users to dissect mitochondrial processes in greater detail.

By leveraging the proposed mitochondrial gene sets, mitology allows for in-depth analysis using transcriptomic data, such as single-transcriptome expression data from RNA sequencing, single-cell sequencing, or spatial transcriptomics.

Mitology is an ongoing project with great potential to advance mitochondrial research. By enabling more detailed insights into mitochondrial contributions to cellular signaling, it provides a valuable tool for exploring mitochondrial biology, disease mechanisms, and therapeutic targets. We are confident that large-scale applications of mitology to publicly available datasets will reveal novel insights, especially in areas where the role of mitochondria has been underexplored due to the limitations of traditional signaling pathway analysis.

As new data and updated annotations become available, we anticipate that mitology will continue to evolve, offering an increasingly comprehensive framework for understanding the molecular complexity of mitochondria.

